# Quantitative visualization of gene expression in *Pseudomonas aeruginosa* aggregates reveals peak expression of alginate in the hypoxic zone

**DOI:** 10.1101/632893

**Authors:** Peter Jorth, Melanie A. Spero, Dianne K. Newman

## Abstract

It is well appreciated that oxygen- and nutrient-limiting gradients characterize microenvironments within chronic infections that foster bacterial tolerance to treatment and the immune response. However, determining how bacteria respond to these microenvironments has been limited by a lack of tools to study bacterial functions at the relevant spatial scales *in situ*. Here we report the application of the hybridization chain reaction (HCR) v3.0 to *Pseudomonas aeruginosa* aggregates as a step towards this end. As proof-of-principle, we visualize the expression of genes needed for the production of alginate (*algD*) and the dissimilatory nitrate reductase (*narG*). Using an inducible bacterial gene expression construct to calibrate the HCR signal, we were able to quantify *algD* and *narG* gene expression across microenvironmental gradients both within single aggregates and within aggregate populations using the Agar Block Biofilm Assay (ABBA). For the ABBA population, alginate gene expression was restricted to hypoxic regions within the environment (~40-200 μM O_2_), as measured by an oxygen microelectrode. Within individual biofilm aggregates, cells proximal to the surface expressed alginate genes to a greater extent than interior cells. Lastly, mucoid biofilms consumed more oxygen than nonmucoid biofilms. These results establish that HCR has a sensitive dynamic range and can be used to resolve subtle differences in gene expression at spatial scales relevant to microbial assemblages. Because HCR v3.0 can be performed on diverse cell types, this methodological advance has the potential to enable quantitative studies of microbial gene expression in diverse contexts, including pathogen behavior in human chronic infections.

**Importance:** The visualization of microbial activities in natural environments is an important goal for numerous studies in microbial ecology, be the environment a sediment, soil, or infected human tissue. Here we report the application of the hybridization chain reaction (HCR) v3.0 to measure microbial gene expression *in situ* at single-cell resolution in aggregate biofilms. Using *Pseudomonas aeruginosa* with a tunable gene expression system, we show that this methodology is quantitative. Leveraging HCR v3.0 to measure gene expression within a *P. aeruginosa* aggregate, we find that bacteria just below the aggregate surface are the primary cells expressing genes that protect the population against antibiotics and the immune system. This observation suggests that therapies targeting bacteria growing with small amounts of oxygen may be most effective against these hard-to-treat infections. More generally, HCR v3.0 has potential for broad application into microbial activities *in situ* at small spatial scales.

## Observation

Despite decades of research that have elucidated mechanisms of bacterial virulence, antibiotic tolerance, and antibiotic resistance, many infections remain impossible to eradicate. Phenotypic heterogeneity likely plays an important role in the failure of drugs and the immune system to clear chronic infections. Chronic *Pseudomonas aeruginosa* lung infections in people with cystic fibrosis (CF) are a prime example. Within individual lobes of the CF lung, genetically antibiotic susceptible and resistant *P. aeruginosa* co-exist (1). This likely affects treatment because resistant bacteria can protect susceptible bacteria when mixed together *in vitro* (2, 3). Likewise, CF lung mucus contains steep oxygen gradients, and anoxic conditions reduce antibiotic susceptibility (4–7). While we know that bacterial genetic diversity and infection site chemical heterogeneity exist, tools to measure bacterial phenotypes *in situ* are lacking. Here we tested the ability of the third generation of the hybridization chain reaction (HCR v3.0) to quantitatively measure gene expression in *P. aeruginosa* in an aggregate model system.

### In situ HCR v3.0 is specific and quantitative for bacterial gene expression

HCR is a fluorescent *in situ* hybridization-like approach that includes a signal amplification step to help visualize low-abundant RNAs (8, 9). We previously used single HCR v2.0 probes to detect bacterial taxa in CF sputum samples (10), and HCR 2.0 was also used by Nikolakakis *et al.* to detect host and bacterial mRNAs in the Hawaiian bobtail squid-*Vibrio fischeri* symbiosis (11). We chose to test HCR v3.0 as a tool to quantify bacterial gene expression *in situ* because of its improved specificity over HCR v2.0. HCR v3.0 requires two paired initiator probes to anneal adjacent to one another on each RNA target before signal amplification occurs, which reduces background signal compared to HCR v2.0 non-specific binding of single initiator probes (Fig. 1A) (8, 9). Therefore, we designed and validated two types of HCR v3.0 probes which could be used to 1) differentiate species and 2) measure gene expression.

**Figure 1.**
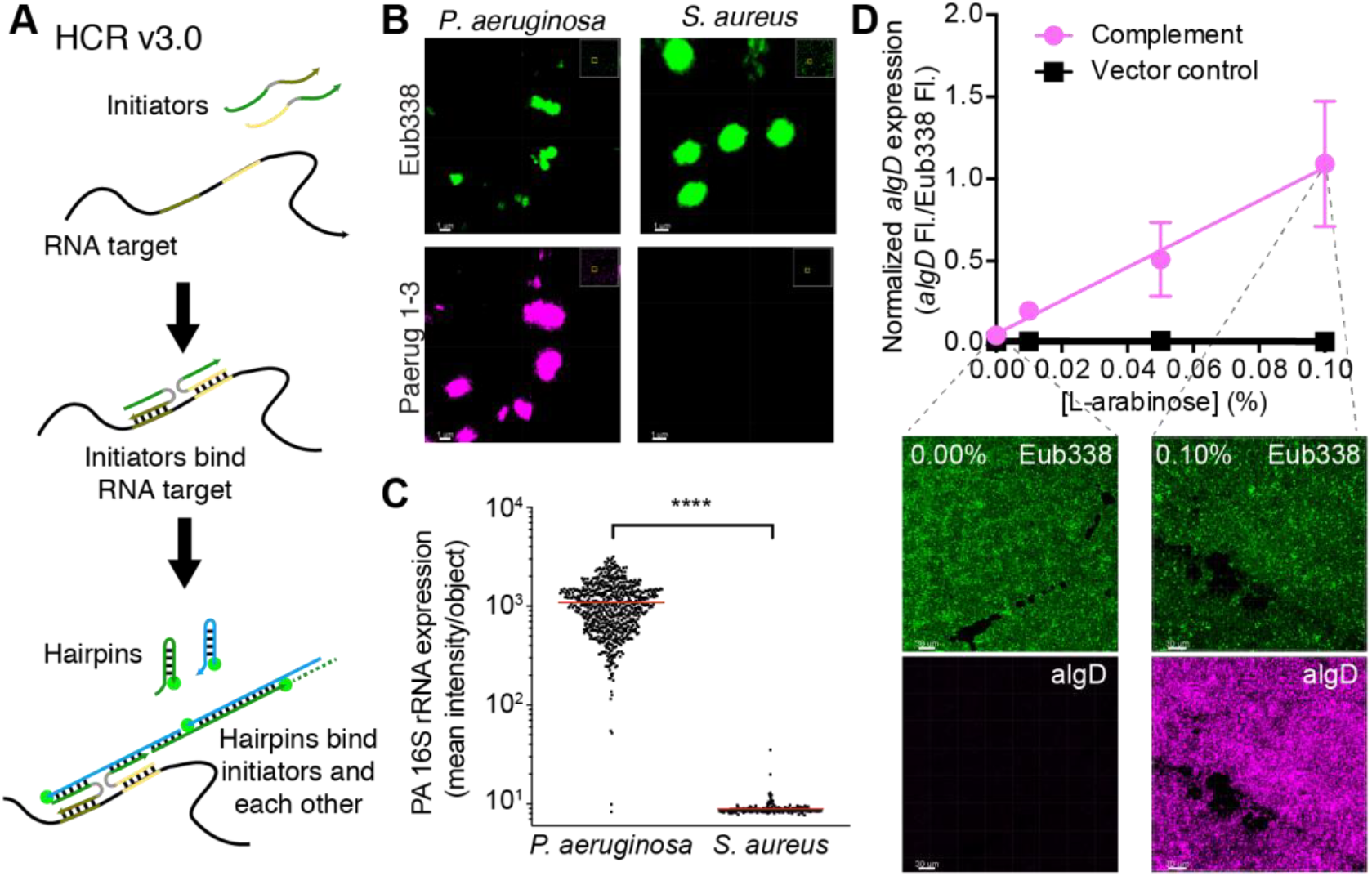
HCR v3.0 analysis is specific and quantitative. **A.** HCR v3.0 utilizes two initiator probes that bind an RNA target followed by two hairpin probes that bind the initiators and each other to generate an amplified fluorescent signal. **B.** HCR v3.0 probes bind to intended targets in bacterial cells. Micrographs show that the Eub338 probe pair (green) binds both *P. aeruginosa* and *S. aureus* rRNA, while the *P. aeruginosa* Paerug1-3 probe pairs mixture (magenta) binds only *P. aeruginosa* rRNA. Scale bars indicate 1 μm. **C.** The *P. aeruginosa* Paerug1-3 probe pairs mixture binds more to individual *P. aeruginosa* cells than *S. aureus* cells (red lines: mean; ****p<0.0001, unpaired t-test). **D.** *algD* mRNA-HCR analysis shows fluorescence increases linearly in the PAO1 Δ*algD* pMQ72::*algD* strain (complement) with increasing arabinose inducer, while the PAO1 Δ*algD* pMQ72 strain (vector control) shows no induction. Representative micrographs for the complement strain with 0% and 0.10% arabinose inducer are shown. Mean normalized fluorescence (*algD* relative to rRNA) is graphed (complement n=3 images per concentration, mean +/− SEM; vector control n=2 images per concentration, mean only). Scale 30 μm. See also Figures S1, S2, and S3.

Using our previous HCR v2.0 probes as a template (10), we designed HCR v3.0 probes to detect 16S rRNA in all eubacteria and *P. aeruginosa* specifically. As expected, the *P. aeruginosa* probes detected only *P. aeruginosa* and not *Staphylococcus aureus*, while the eubacterial probe detected both organisms (Fig. 1B-C and S1). When only one initiator probe from each pair was used, no fluorescence was observed, as anticipated (Fig. S2). Thus, HCR v3.0 probes were highly specific for the intended bacteria.

To test the ability of HCR v3.0 to quantify bacterial gene expression, we designed probes to detect *P. aeruginosa algD* mRNA, and we cloned *algD* into the arabinose-inducible expression plasmid pMQ72 in a *P. aeruginosa* Δ*algD* mutant (12, 13). mRNA-HCR analysis was highly quantitative: we observed a linear relationship between the concentration of the inducer (*i.e.* expression level) and HCR signal in the complemented strain, while the empty vector control strain produced no signal (Fig. 1D and S3). This demonstrated that mRNA-HCR can quantify bacterial gene expression *in situ*.

### mRNA-HCR reveals alginate gene expression in hypoxic zones of P. aeruginosa aggregates

As a case study, we chose to measure *P. aeruginosa* alginate (*algD*) and nitrate reductase (*narG*) gene expression in aggregates formed by a mucoid (FRD1) and nonmucoid strain (PA14). This approach was chosen for several reasons. First, measuring *algD* expression *in situ* is of interest because alginate is overproduced by mucoid strains in CF lung infections (14, 15), and mucoid strains are associated with worsened lung function (16). Second, as a technical control, the *algD* gene should be more highly expressed in the mucoid than in nonmucoid strain and produce a stronger HCR signal (14). Third, previous research suggests that alginate may be expressed in hypoxic and anoxic conditions (7, 17-19), yet the precise location of alginate gene expression in aggregate biofilms has yet to be determined. Therefore, we could also quantify *algD* expression relative to *narG*, a gene induced under hypoxic and anoxic conditions (17, 20), which would help determine where *algD* is expressed in aggregates relative to environmental oxygen availability.

Using the Agar Block Biofilm Assay (ABBA) (20), we grew mucoid and nonmucoid aggregates suspended in an agar medium and measured *narG* and *algD* gene expression with mRNA- HCR. As expected, the mucoid strain expressed *algD* more highly than the nonmucoid strain (Fig. 2A-C,E). Spatially, *algD* expression was highest in the zones within the first 200 μm below the air-agar interface (Fig 2A-C,E). Interestingly, *narG* was also expressed more highly in the mucoid than nonmucoid strain (Fig 2D) and was expressed more evenly in aggregates at varying depths below the agar surface (Fig. 2A-D). Analysis of individual aggregates in the ABBA experiments showed an intriguing ring-like pattern of 16S rRNA, *algD*, and *narG* gene expression. Within individual aggregates, *algD* expression was detected in cells ~5-15 μm below the aggregate surface but was not detected in the innermost cells within ~10 μm of the aggregate center (Fig. 2F-G). In contrast, the innermost cells highly expressed *narG*, but cells within ~0-10 μm of the aggregate surface did not express *narG* (Fig. 2F-G). This led us to hypothesize that *algD* was being expressed by cells experiencing hypoxia just below the aggregate surface and not by cells in the innermost, presumably anoxic, regions of the aggregates.

**Figure 2.**
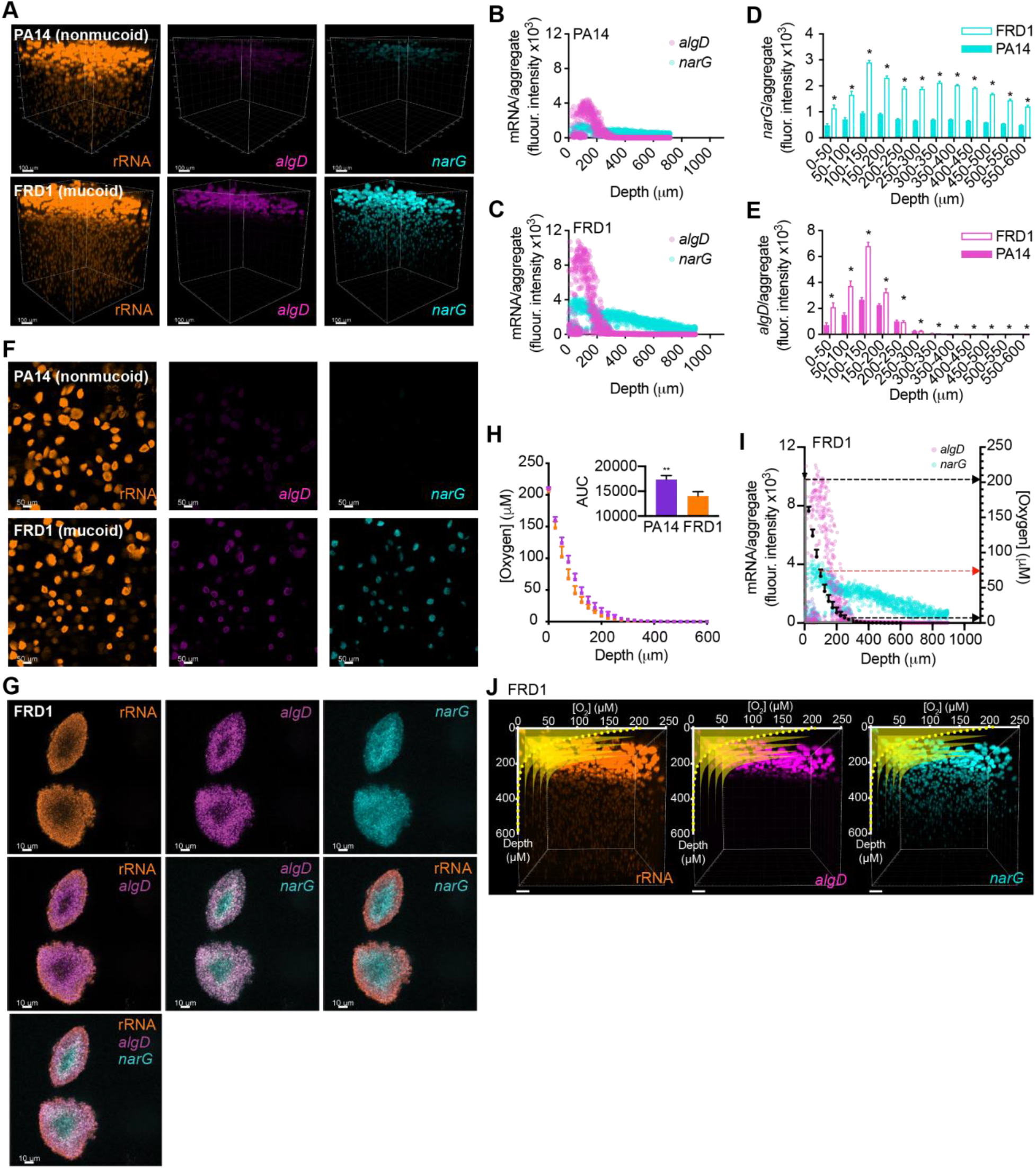
Alginate gene expression is highest in hypoxic regions of *P. aeruginosa* aggregates. **A.** 3D fluorescence micrographs of nonmucoid PA14 and mucoid FRD1 ABBA samples probed with the Eub338 (rRNA), *algD*, and *narG* HCR v3.0 probes. Scale bars: 100 μm. **B-C.** Mean *algD* and *narG* HCR signals per individual aggregate in nonmucoid (**B**) and mucoid (**C**) strains. **D-E.** Mean *algD* (**D**) and *narG* (**E**) HCR signals per ABBA aggregate biofilm at different binned depths below the air-agar interface in each sample (50 μm bins; mean +/− SEM; *p<0.05, unpaired two-tailed t-test; nd: not detected). **F.** 2D micrographs of nonmucoid and mucoid ABBA samples probed with the rRNA, *algD*, and *narG* HCR probes. Images correspond to single Z-slices 99 μm below the air-agar interface. Scale bars: 50 μm. **G.** 2D micrograph of mucoid ABBA aggregates probed with the rRNA, *algD*, and *narG* HCR probes. Overlays show that nitrate reductase is expressed by interior bacterial cells, while *algD* is expressed by bacterial cells just below the aggregate surface. Each image corresponds to the same Z-slice with different probes shown. Scale bars: 10 μm. **H.** Oxygen profiles in nonmucoid and mucoid ABBA samples. Mean oxygen concentrations at 25 μm intervals from the air-agar interface to 600 μm below (n=3) are indicated. Inset bar graph indicates area under the curve (AUC) for each scatter plot (**p<0.005, unpaired two-tailed t-test). **I.** Mean *algD* and *narG* expression per mucoid ABBA aggregate (left y-axis) plotted with mean oxygen concentrations measured (right y-axis). Red arrow indicates oxygen concentration at which peak *algD* expression was detected, black arrows indicate minimum and maximum oxygen concentrations at which *algD* expression was detected. In **H&I**, error bars indicate SEM for the oxygen concentrations. **J.** Expression of *algD* is restricted to hypoxic regions, while *narG* is detected in hypoxic, and anoxic regions. Oxygen profiles (yellow) overlay 3D micrographs showing rRNA, *algD*, and *narG* HCR signals in mucoid ABBA samples. Oxygen profiles are plotted multiple times using perspective at different xz-planes along the y-axis. In **A-G,&I-J** data are shown from a representative ABBA experiment. Results from a replicate experiment are shown in Figure S4.

To test where cells were expressing *algD* relative to oxygen availability, we used a microelectrode to measure oxygen concentrations from 0-600 µm below the agar surface in mucoid and nonmucoid ABBA experiments. Unexpectedly, the mucoid strain showed a modest increase in its oxygen consumption rate compared to the nonmucoid strain (Fig. 2H). However, as we predicted, the mucoid strain expressed *algD* highest in hypoxic regions (5-200 μM oxygen) of the agar, from 0-350 μm below the agar surface and peaking at ~75 μM oxygen (Fig. 2I-J). In regions with less than 5 μM oxygen, *algD* expression plummeted to <1% of the maximum value detected (Fig. 2I-J). This was surprising because in planktonic cultures we found that anoxia most strongly induced *algD* expression compared to oxic and hypoxic conditions (Fig. S5), similar to previous research (18). Thus, alginate gene expression patterns differ between planktonic and aggregate cells: in aggregate cells, *algD* expression is greatest under hypoxic rather than anoxic conditions.

## Conclusion

Altogether, these experiments demonstrate the utility of HCR v3.0 for quantitatively measuring bacterial gene expression *in situ* at spatial scales relevant to microbial assemblages. Going forward, it will be exciting to combine mRNA-HCR with tissue clearing methods such as MiPACT (10) to determine whether the expression patterns observed in these *in vitro* studies similarly characterize aggregate populations of pathogens *in vivo*. Direct insight into how pathogen physiology develops in infected tissues, or any other context where spatial observation of microbial activities is important, promises to yield insights that will facilitate more effective control of these communities. Many applications of HCR v3.0 can be envisioned, such as using this visualization tool to analyze microbes after therapeutic interventions to identify bacterial subpopulations that either resist or succumb to treatment. Ultimately, identifying the subpopulations that survive a specific perturbation can be used to guide the development and implementation of future therapeutics.

## Methods

Bacterial strains were routinely grown in Luria Bertani broth and agar. Bacterial cloning, ABBA experiments, HCR analyses, and oxygen measurements were performed as described previously (10, 12, 21–25). For experimental details see Supplemental Methods and Tables including probe sequences (Table S1), bacterial strains (Table S2), and primers (Table S3).

## Acknowledgements

We would like to thank Will DePas, Ruth Lee, Niles Pierce and the Programmable Molecular Technology Center at the Caltech Beckman Institute for technical assistance and advice. Confocal microscopy was performed in the Caltech Biological Imaging Facility at the Caltech Beckman Institute, which is supported by the Arnold and Mabel Beckman Foundation. Grants to DKN from the Army Research Office (W911NF-17-1-0024) and National Institutes of Health (1R01AI127850-01A1) supported this research. PJ was supported by postdoctoral fellowships from the Cystic Fibrosis Foundation (JORTH14F0 and JORTH17F5). MAS was supported by a gift from the Doren Family Foundation.

